# Seasonal temperature variation influences climate suitability for dengue, chikungunya, and Zika transmission

**DOI:** 10.1101/230383

**Authors:** John H. Huber, Marissa L. Childs, Jamie M. Caldwell, Erin A. Mordecai

## Abstract

Dengue, chikungunya, and Zika virus epidemics transmitted by *Aedes aegypti* mosquitoes have recently (re)emerged and spread throughout the Americas, Southeast Asia, the Pacific Islands, and elsewhere. Understanding how environmental conditions affect epidemic dynamics is critical for predicting and responding to the geographic and seasonal spread of disease. Specifically, we lack a mechanistic understanding of how seasonal variation in temperature affects epidemic magnitude and duration. Here, we develop a dynamic disease transmission model for dengue virus and *Aedes aegypti* mosquitoes that integrates mechanistic, empirically parameterized, and independently validated mosquito and virus trait thermal responses under seasonally varying temperatures. We examine the influence of seasonal temperature mean, variation, and temperature at the start of the epidemic on disease dynamics. We find that at both constant and seasonally varying temperatures, warmer temperatures at the start of epidemics promote more rapid epidemics due to faster burnout of the susceptible population. By contrast, intermediate temperatures (24-25°C) at epidemic onset produced the largest epidemics in both constant and seasonally varying temperature regimes. When seasonal temperature variation was low, 25-35°C annual average temperatures produced the largest epidemics, but this range shifted to cooler temperatures as seasonal temperature variation increased (analogous to previous results for diurnal temperature variation). Tropical and sub-tropical cities such as Rio de Janeiro, Fortaleza, and Salvador, Brazil; Cali, Cartagena, and Barranquilla, Colombia; Delhi, India; Guangzhou, China; and Manila, Philippines have mean annual temperatures and seasonal temperature ranges that produced the largest epidemics. However, more temperate cities like Shanghai, China had high epidemic suitability because large seasonal variation offset moderate annual average temperatures. By accounting for seasonal variation in temperature, the model provides a baseline for mechanistically understanding environmental suitability for virus transmission by *Aedes aegypti*. Overlaying the impact of human activities and socioeconomic factors onto this mechanistic temperature-dependent framework is critical for understanding likelihood and magnitude of outbreaks.

**Non-Technical Summary:** Mosquito-borne viruses like dengue, Zika, and chikungunya have recently caused large epidemics that are partly driven by temperature. Using a mathematical model built from laboratory experimental data for *Aedes aegypti* mosquitoes and dengue virus, we examine the impact of variation in seasonal temperature regimes on epidemic size and duration. At constant temperatures, both low and high temperatures (20°C and 35°C) produce small epidemics, while intermediate temperatures like 25°C and 30°C produce much larger epidemics. In seasonally varying temperature environments, epidemics peak more rapidly at higher starting temperatures, while intermediate starting temperatures produce the largest epidemics. Seasonal mean temperatures of 25–35°C are most suitable for large epidemics when seasonality is low, but in more variable seasonal environments epidemic suitability peaks at lower annual average temperatures. Tropical and sub-tropical cities have the highest temperature suitability for epidemics, but more temperate cities with high seasonal variation also have the potential for very large epidemics.

## Introduction

Over the last 30–40 years, arboviral outbreaks have dominated the public health landscape globally [1]. These viruses, most notably dengue (DENV), chikungunya (CHIKV), and Zika (ZIKV), can cause symptoms ranging from rash, arthralgia, and fever to hemorrhagic fever (DENV), long-term arthritis (CHIKV), Guillain-Barré syndrome and microcephaly (ZIKV) [2–4]. DENV, which historically spread worldwide along shipping routes [5], places 3.97 billion individuals at risk worldwide [6] and causes an estimated 390 million cases annually, including 96 million symptomatic cases [7]. CHIKV was introduced into the Americas in December 2013 after an outbreak in St. Martin Island [8]. Since then, autochthonous transmission has been reported in 45 countries [9], and 1.3 billion people worldwide are at risk of contracting CHIKV [10]. More recently, the ZIKV epidemic in the Americas captured global attention after the World Health Organization (WHO) designated it a Public Health Emergency of International Concern in February 2016 in response to its association with neurological disorders. Following the first reported case in Brazil in May 2015, ZIKV has spread to 48 countries and territories where it is transmitted autochthonously [11]. Because DENV, CHIKV, and ZIKV are mostly transmitted by *Aedes aegypti* mosquitoes, they may have similar geographic distributions and risk factors.

Informed public health decisions to limit the spread and magnitude of these arboviral epidemics depend on a robust understanding of transmission dynamics. One mechanistic modeling framework, the Susceptible – Infected – Recovered (SIR) model, has been implemented successfully to model the dynamics of outbreaks of influenza, measles, and vector-borne diseases such as CHIKV and ZIKV [12–14]. This approach tracks virus population dynamics by compartmentalizing individuals by their state in an epidemic (i.e., Susceptible (S), Infected (I), Recovered (R)). This framework can be extended to include additional compartments, such as a latency stage, or to incorporate the dynamics of the mosquito population for vector transmission.

Arbovirus dynamics are strikingly seasonal and geographically restricted to relatively warm climates [6,7]. This arises because several life history traits of the mosquitoes that transmit DENV, CHIKV, and ZIKV are strongly influenced by temperature and seasonality [15–22]. For simplicity, many existing models assume static life history traits [14], and those that address seasonal forcing tend to incorporate sinusoidal variation as a single transmission parameter, *β* [23]. Treating seasonal temperature variation as a sinusoidal forcing function on the transmission parameter implies a monotonic relationship between temperature and transmission, such that transmission is maximized at high temperatures and decreases at low temperatures. However, decades of experimental work have demonstrated strongly nonlinear (often unimodal) relationships between mosquito and pathogen traits and temperature that are not well captured in a single sinusoidal forcing function [24]. Efforts by Yang et al. [25,26] addressed the need to include seasonal variation by adopting an SEI-SEIR compartmental framework with time-varying entomological parameters and fitting the model to DENV incidence data in Campinas, Brazil. Other previous work has integrated the effects of temperature on mosquito and parasite traits into temperature-dependent transmission models for DENV, CHIKV, and/or ZIKV, and revealing a strong, nonlinear influence of temperature with peak transmission between 29 – 35 °C [27–34]. However, we do not yet have a mechanistic estimate for the relationship between seasonal temperature regimes and transmission potential, incorporating the full suite of transmission-relevant, nonlinear thermal responses of mosquito and parasite traits.

Here, we expand on previous work with three main advances: (1) we incorporate the full suite of empirically-derived, unimodal thermal responses for all known transmission-relevant mosquito and parasite traits; (2) we examine the influence of seasonal temperature mean and variation (in contrast to constant temperatures or daily temperature variation); and (3) we use a dynamic transmission framework to explore the impact of different seasonal temperature regimes on the epidemiologically-relevant outcomes of epidemic size, duration, and peak incidence (in contrast to R0, or vectorial capacity, which are difficult to measure directly). To do so, we incorporate previously estimated and independently validated thermal response functions for all vector and parasite traits [24] into a dynamic SEI-SEIR model [25,26]. We explore field-relevant temperature regimes by simulating epidemics across temperature means (10 – 38°C) and seasonal ranges (0 – 17°C) from across the predicted suitable range for transmission. Specifically, we use the model to ask: (1) How does final epidemic size vary across constant temperatures? (2) Under seasonally varying temperatures, how does the temperature at the start of the epidemic affect the final epidemic size and duration? (3) How do temperature mean and seasonal range interact to determine epidemic size? (4) Which geographic locations have high epidemic suitability based on climate?

## Methods

### Model

#### Model Framework

We adopted an SEI-SEIR compartmental modeling framework to simulate arboviral transmission by the *Aedes aegypti* vector (Fig. 1). We introduced temperature-dependence into the model by using fitted thermal response curves for the mosquito life history traits provided by Mordecai et al. [24]. The full model is:

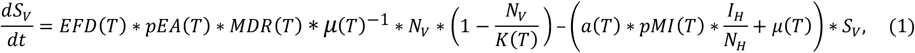

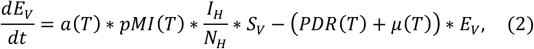

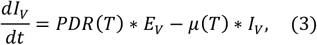

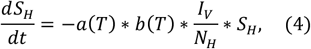

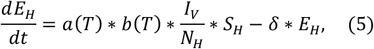

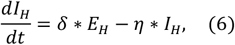

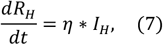

**Fig 1.**
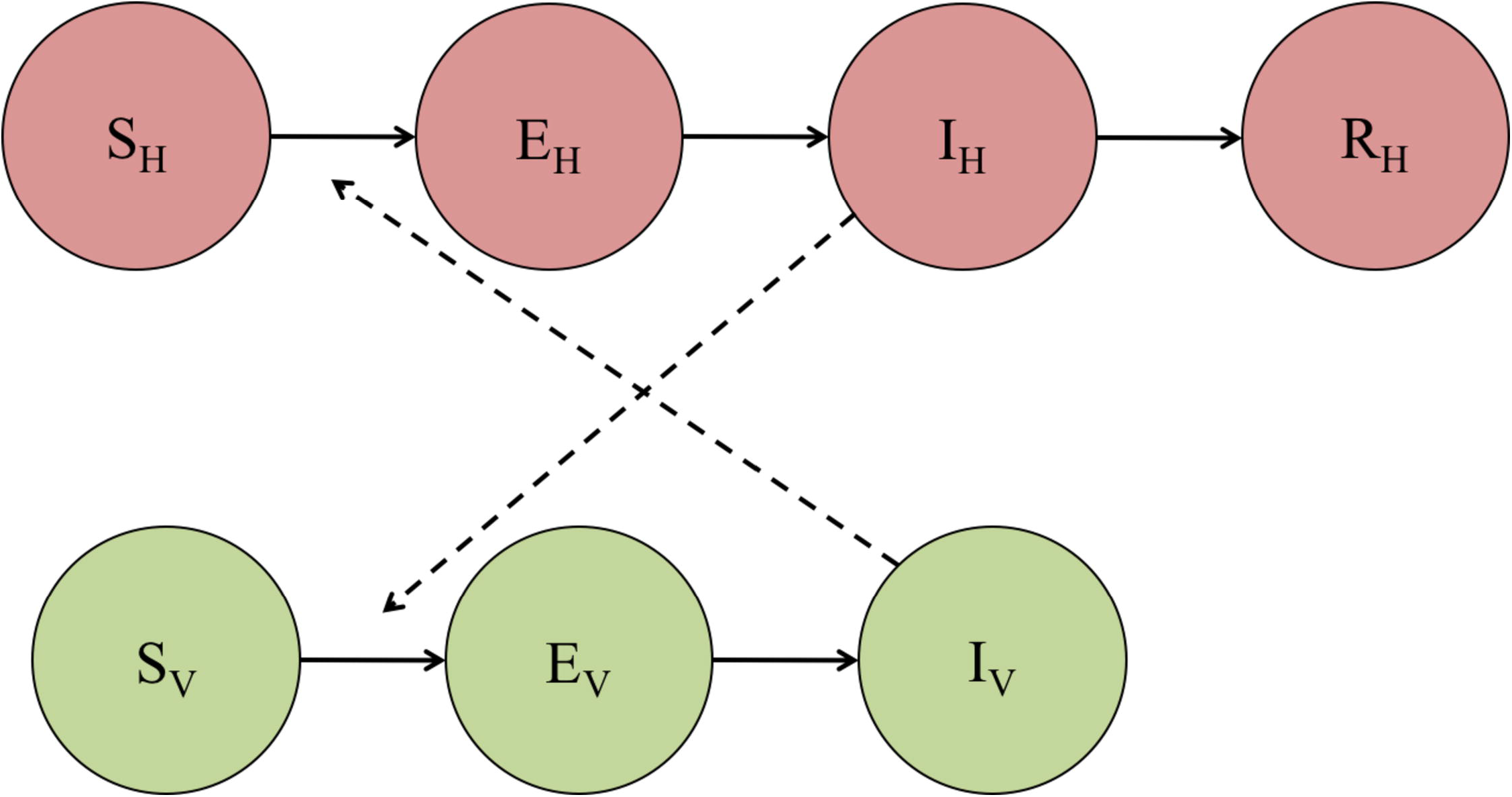
Compartmental model of transmission. *S_H_, E_H_, I_H_*, and *R_H_* represent the susceptible, exposed (or latent), infectious, and recovered segments of the human population, respectively. Likewise, *S_V_, E_V_*, and *I_V_* represent the susceptible, exposed (or latent), and infectious segments of the mosquito population. Solid arrows signify the directionality of transition from one compartment to the next, and dashed arrows indicate the directionality of transmission.

The SEI portion of the model describes the vector population, where *S_V_* represents the number of susceptible mosquitoes, *E_V_* is the number of mosquitoes in the latency stage, and *I_V_* is the number of infectious mosquitoes. We assumed that *Aedes aegypti* mosquitoes remain infectious until they die. In equations 1-3, (*T*) indicates temperature-dependent functions, *EFD*(*T*) is the number of eggs laid per female per day, *pEA*(*T*) is the probability of mosquito egg-to-adult survival, *MDR*(*T*) is the mosquito egg-to-adult development rate, *N_V_* is the total mosquito population at time *t* (i.e., *S_V_* + *E_V_* + *I_V_*), *K*(*T*) is the carrying capacity for the mosquito population, *a*(*T*) is the per mosquito biting rate, *pMI*(*T*) is the probability of mosquito infection per bite on an infectious host, *μ*(*T*) is the adult mosquito mortality rate, and *PDR*(*T*) is the parasite development rate. Each life history and pathogen transmission trait of the *Aedes aegypti* mosquito is a unimodal, temperature-dependent function fit from experimental laboratory data in previous work [15–22,24] (Table 1; Appendix; “Functional Forms of Life History Traits”).

**Table 1.** Fitted thermal responses for *Aedes aegypti* life history traits. Traits were fit to a Brière 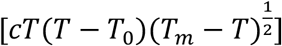 or a quadratic [*c*(*T* – *T_m_*)(*T* – *T*_0_)] function where *T* represents temperature. *T*_0_ and *T_m_* are the critical thermal minimum and maximum, respectively, and *c* is the rate constant. Thermal responses were fit by [24].

| Trait | Definition | Function | Fitted Parameters |  |  |
| --- | --- | --- | --- | --- | --- |
| $a$ | Biting rate (day<sup>-1</sup>) | Brière | $c = 2.02\text{e-}04$ | $T_{min} = 13.35$ | $T_{max} = 40.08$ |
| $EFD$ | Eggs laid per female per day | Brière | $c = 8.56\text{e-}03$ | $T_{min} = 14.58$ | $T_{max} = 34.61$ |
| $pEA$ | Probability of mosquito egg-to-adult survival | Quadratic | $c = -5.99\text{e-}03$ | $T_{min} = 13.56$ | $T_{max} = 38.29$ |
| $MDR$ | Mosquito egg-to-adult development rate (day<sup>-1</sup>) | Brière | $c = 7.86\text{e-}05$ | $T_{min} = 11.36$ | $T_{max} = 39.17$ |
| $lf$ | Adult mosquito lifespan (days) | Quadratic | $c = -1.48\text{e-}01$ | $T_{min} = 9.16$ | $T_{max} = 37.73$ |
| $b$ | Probability of mosquito infectiousness | Brière | $c = 8.49\text{e-}04$ | $T_{min} = 17.05$ | $T_{max} = 35.83$ |
| $pMI$ | Probability of mosquito infection | Brière | $c = 4.91\text{e-}04$ | $T_{min} = 12.22$ | $T_{max} = 37.46$ |
| $PDR$ | Virus extrinsic incubation rate (day<sup>-1</sup>) | Brière | $c = 6.65\text{e-}05$ | $T_{min} = 10.68$ | $T_{max} = 45.90$ |

The SEIR portion of the model describes the human population, where *S_H_* represents the number of susceptible individuals, *E_H_* the number of latent (or exposed) individuals, *I_H_* the number of infectious individuals, and *R_H_* the number of recovered individuals. We assumed a static population size, *N_H_*, that was neither subject to births nor deaths because the human lifespan far exceeds the duration of an epidemic. Further, we binned asymptomatic and symptomatic individuals into a single infectious class since asymptomatic infections have been shown to transmit DENV [35] and exhibit similar viremic profiles as symptomatic patients in CHIKV [36]. Based on previous arboviral outbreaks [37,38], we assumed that an infection conferred long-term immunity to an individual. Thus, a previously infectious individual entering the recovered class is protected from subsequent re-infection for the remainder of the epidemic. In the case of dengue, where there are four unique serotypes, we consider single-season epidemics of a single serotype. In equations 4-7, *b*(*T*) is the probability of human infection per bite by an infectious mosquito (Table 1), δ^−1^ is the intrinsic incubation period, and η^−1^ is the human infectivity period. Since human components of the transmission cycle are not seasonal, we used constants of 5.9 days for the intrinsic incubation period, 1/δ, and 5.0 days for the infectious period, 1/η [14]. All temperature-independent parameter values are given in Table 2.

**Table 2.**
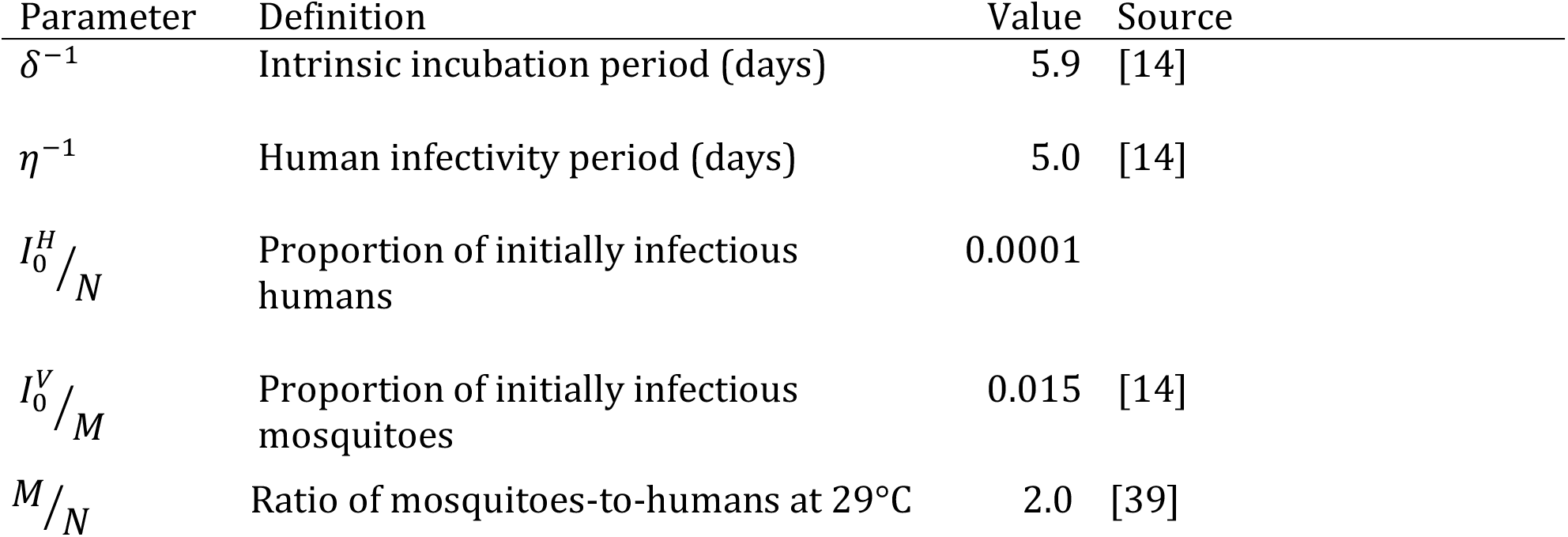
Values of temperature-independent parameters used in the model, and their sources.

Since the lifespan of an adult mosquito is short relative to the timespan of an epidemic, we allowed mosquito birth and death rates to drive population dynamics. Additionally, the birth rate of susceptible mosquitoes was regulated by a temperature-dependent carrying capacity, *K* (equation 8), which we modeled as a modified Arrhenius equation that is a unimodal function of temperature [40]:

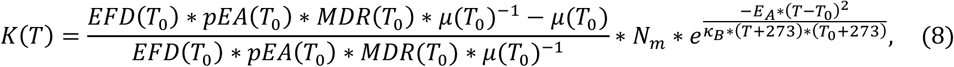

Here, *T*_0_ is defined as the reference temperature (i.e., the temperature at which the carrying capacity is greatest) in Celsius, *N_m_* is the maximum carrying capacity, and *κ_B_* is Boltzmann constant (8.617 × 10^−5^ eV/K). *EFD* is the number of eggs laid per female per day, *pEA* is the probability of egg-to-adult mosquito survival, *MDR* is the mosquito egg-to-adult development rate, and *μ* is the adult mosquito mortality rate. We calculated these values for the reference temperature. *E_A_* is the activation energy, which we set to 0.5 and represents the temperature dependence of the carrying capacity, a conservative estimate as we lacked sufficient data on estimates of the carrying capacity of *Aedes aegypti* and its underlying temperature dependence. To convert from Celsius to Kelvin, we incremented the temperature *T* and the reference temperature *T*_0_ by 273. Equation (8) was adopted from [40] and modified to allow the distribution to be unimodal. We set the reference temperature, *T*_0_, to 29°C, which is consistent with optimal temperatures for *Aedes aegypti* transmission [24,29].

We included a temperature-dependent carrying capacity in the model to constrain the growth of the mosquito population. As described in the Appendix, all simulations begin with the mosquito population at its (temperature-dependent) carrying capacity. As the temperature changes seasonally, the mosquito population does not necessarily remain at carrying capacity if one or more of the life history traits that determine the production of new mosquitoes in equation (1)—*EFD, pEA*, and *MDR*—is equal to zero. This occurs below 14.58°C (the highest *Tmin* of *EFD, pEA*, and *MDR*) or above 34.61°C (the lowest *Tmax* of *EFD, pEA*, and *MDR*).

It should be noted that the transmission parameters are only related to the current temperature at each time point in the simulation. Time lags for each life history trait were not explicitly built into the model.

#### Seasonal Forcing

To address seasonality in the model, we allowed temperature to vary over time. We modeled temperature as a sinusoidal curve with a period of 365 days of the form:

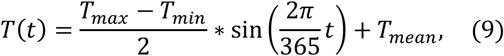

Here, *T_max_, T_mean_*, and *T_min_* represent the average monthly maximum, mean, and minimum temperatures across a calendar year, respectively, and *t* is measured in days. By modeling temperature as a function of time, we allowed the life history traits of the *Aedes aegypti* vector to vary across time for the duration of the epidemic. In the absence of a specific focal location we modeled seasonal temperature as a sinusoidal function for simplicity.

### Data

#### Life History Traits

To incorporate seasonal forcing into the compartmental modeling framework, we used fitted mechanistic thermal response curves [24]. Mordecai et al. [24] examined published data on thermal responses for life history traits of the *Aedes aegypti* vector and DENV and adopted a Bayesian approach for fitting quadratic (*Q*(*T*); Eq. 10) or Brière (*B*(*T*); Eq. 11) curves (see Appendix for details).

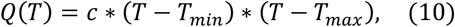

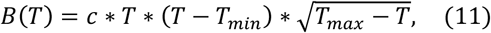

Here, *c* is a rate constant, *T_min_* is the critical temperature minimum, and *T_max_* is the critical temperature maximum (Table 1). Following Mordecai et al. [24], we assumed values above the critical thermal maxima and below the minima were equal to zero.

Mordecai et al. [24] fit the thermal response for adult mosquito lifespan (Table 1), the inverse of the adult mosquito mortality rate (*μ*, in days^−1^), used in our model. We set the mortality rate at temperatures outside the critical thermal minimum and maximum to 24 days^−1^ (i.e., mosquitoes survive for one hour at temperatures outside of the Tmin to Tmax range).

#### Historical Weather Data

To identify areas of epidemic suitability across the globe, we extracted monthly mean temperatures for 2016 from Weather Underground (wunderground.com) for twenty different cities (Table 3). For each city, we calculated the mean, minimum, and maximum from the average monthly mean temperatures, to estimate temperature seasonality. This provided a range of the average monthly temperature over the span of a calendar year. We chose this time period because it provided the most recent full calendar year to demonstrate seasonal variation in temperature.

**Table 3.** Temperature regimes for major cities during the 2016 calendar year. Monthly mean temperatures during 2016 were extracted from Weather Underground.

| City | Annual Mean Temperature<br>(°C) | Annual Temperature<br>Amplitude (°C) |
| --- | --- | --- |
| Buenos Aires, Argentina | 16.5 | 8.0 |
| Sao Paulo, Brazil | 20.6 | 5.0 |
| Rio de Janeiro, Brazil | 24.3 | 4.0 |
| Salvador, Brazil | 26.3 | 2.0 |
| Fortaleza, Brazil | 27.8 | 0.50 |
| Belo Horizonte, Brazil | 21.9 | 3.0 |
| Recife, Brazil | 27.2 | 1.5 |
| Shanghai, China | 17.6 | 12.5 |
| Beijing, China | 12.8 | 16 |
| Guangzhou, China | 22.9 | 8.0 |
| Bogotá, Colombia | 14.7 | 1.0 |
| Medellin, Colombia | 17.9 | 1.0 |
| Cali, Colombia | 25.1 | 1.5 |
| Barranquilla, Colombia | 28.8 | 1.0 |
| Cartagena, Colombia | 28.6 | 1.0 |
| Delhi, India | 26.3 | 9.5 |
| Tokyo, Japan | 17.0 | 10.5 |
| Kobe, Japan | 17.4 | 11 |
| Manila, Philippines | 29.0 | 1.5 |
| New York, USA | 13.8 | 12 |

**Table 3.** Estimates of epidemic suitability for major cities. Epidemic suitability was calculated as the proportion of the population that became infected in simulations run with 0, 20, 40, 60, or 80% initial population immunity. Temperature at simulation onset was set to the mean of the temperature regime. Each city was simulated with its respective temperature regime from the 2016 calendar year.

| Epidemic Suitability |  |  |  |  |  |
| --- | --- | --- | --- | --- | --- |
| City | 0% Immunity | 20% Immunity | 40% Immunity | 60% Immunity | 80% Immunity |
| Buenos Aires, Argentina | 0.03656 | 0.02169 | 0.01203 | 0.005975 | 0.002295 |
| Sao Paulo, Brazil | 0.6056 | 0.3386 | 0.1518 | 0.05351 | 0.01385 |
| Rio de Janeiro, Brazil | 0.9984 | 0.7962 | 0.5891 | 0.3618 | 0.09862 |
| Salvador, Brazil | 0.9990 | 0.7976 | 0.5937 | 0.3804 | 0.1335 |
| Fortaleza, Brazil | 0.9993 | 0.7982 | 0.5953 | 0.3861 | 0.1535 |
| Belo Horizonte, Brazil | 0.5909 | 0.3344 | 0.1544 | 0.05771 | 0.01633 |
| Recife, Brazil | 0.9994 | 0.7985 | 0.5959 | 0.3871 | 0.1517 |
| Shanghai, China | 0.9966 | 0.7878 | 0.5507 | 0.2484 | 0.03456 |
| Beijing, China | 0.5268 | 0.2526 | 0.09058 | 0.02298 | 0.003587 |
| Guangzhou, China | 0.9996 | 0.7989 | 0.5965 | 0.3848 | 0.1254 |
| Bogotá,<br>Colombia | 0.0001000 | 0.0001000 | 0.0001000 | 0.0001000 | 0.0001000 |
| Medellin,<br>Colombia | 0.002544 | 0.002048 | 0.001556 | 0.001068 | 0.0005820 |
| Cali,<br>Colombia | 0.9909 | 0.7822 | 0.5617 | 0.3122 | 0.07217 |
| Barranquilla,<br>Colombia | 0.9997 | 0.7993 | 0.5979 | 0.3928 | 0.1703 |
| Cartagena,<br>Colombia | 0.9997 | 0.7993 | 0.5978 | 0.3923 | 0.1688 |
| Delhi, India | 0.9537 | 0.7215 | 0.4759 | 0.2388 | 0.06803 |
| Tokyo, Japan | 0.7269 | 0.4149 | 0.1758 | 0.05159 | 0.009489 |
| Kobe, Japan | 0.9435 | 0.6669 | 0.3522 | 0.1090 | 0.01632 |
| Manila,<br>Philippines | 0.9998 | 0.7994 | 0.5981 | 0.3933 | 0.1720 |
| New York,<br>USA | 0.04088 | 0.02159 | 0.01041 | 0.004390 | 0.001425 |

### Variability in Epidemic Dynamics with Constant Temperature

We first examined how epidemic dynamics varied across different constant temperatures. Here, we did not introduce seasonal forcing into the model but rather assumed static life history traits for *Aedes aegypti* for the simulation period. We simulated the model under default starting conditions (see Appendix) at four different constant temperatures: 20°C, 25°C, 30°C, and 35°C. These temperatures were chosen to span the range of temperatures at which arbovirus transmission is likely to be possible [24].

### Variability in Epidemic Dynamics with Starting Temperature

Using the model that included seasonal variation in temperature, we examined how the dynamics of an epidemic varied due to the temperature at which the epidemic began, under two temperature regimes. First, we set *T_max_* = 40.0°C, *T_mean_* = 25.0°C, and *T_min_* = 10.0°C in the time-varying seasonal temperature model under default parameters (see Appendix) and varied the temperature at the start of the epidemic from 10.0°C to 40.0°C in increments of 0.1°C. We examined the response of final epidemic size, epidemic length, and maximum instantaneous number of infected individuals. We then repeated this process for a regime with a lower magnitude of seasonal temperature variation: *T_max_* = 30.0°C, *T_mean_* = 25.0°C, and *T_min_* = 20.0°C. By comparing these temperature regimes, we can examine how epidemics respond to starting temperatures that are outside the range of plausible temperatures of arbovirus transmission (regime 1) versus restricted to the plausible temperatures for transmission (regime 2) [24].

### Seasonal Variability of Final Epidemic Size

Using the compartmental modeling framework with the default starting conditions, we examined the variation in final epidemic size as a result of seasonal forcing. To do so, we simulated over a wide range of temperature mean and seasonal variance regimes. The mean annual temperature varied from 10.0°C to 38.0°C in increments of 0.1°C, while the seasonal variation about the mean (i.e., 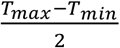) ranged from 0.0°C to 17.0°C in increments of 0.1°C. Many of these temperature regimes are unlikely to be observed empirically. However, the simulated temperature regimes spanned the full range of feasible temperature conditions. We recorded the final epidemic size, measured as the number of individuals in the recovered compartment at the end of the simulation, for each unique combination of mean annual temperature and seasonal variation. In addition, we examined the effect of epidemic starting temperature on final epidemic size across the same seasonal temperature regimes. We ran the model under default starting conditions, but allowed the starting temperature to equal *T_min_, T_mean_*, or *T_max_*.

To observe the interaction of population immunity with the seasonal temperature regime, we simulated the model assuming that 0, 20, 40, 60, or 80% of the population was initially immune. Each simulation began with the introduction of the infected individual occurring at the mean seasonal temperature.

We then compared simulated climate regimes with actual climates in major cities, to measure relative epidemic suitability of the following cities: São Paulo, Brazil; Rio de Janeiro, Brazil; Salvador, Brazil; Fortaleza, Brazil; Belo Horizonte, Brazil; Recife, Brazil; Bogotá, Colombia; Medellín, Colombia; Cali, Colombia; Barranquilla, Colombia; Cartagena, Colombia; Tokyo, Japan; Delhi, India; Manila, Philippines; Shanghai, China; Beijing, China; New York City, USA; Guangzhou, China; Kobe, Japan; and Buenos Aires, Argentina, given 0, 20, 40, 60, and 80% population immunity. These cities were chosen because they represent some of the most populous urban areas across South America and throughout the world.

### Model Sensitivity and Uncertainty Analysis

To characterize uncertainty in the model, we sampled 50 joint posterior estimates for *c*, *T_min_*, and *T_max_* for each life history trait provided by Mordecai et al. [24]. We examined the variability in epidemic dynamics with starting temperatures under each parameterization and report the 95% credible interval for the epidemiological indices. We similarly characterize uncertainty in our estimates of the final epidemic size as a function of the seasonal temperature regime by simulating under each parameterization and reporting the 95% credible interval.

## Results

### Variability in Epidemic Dynamics with Constant Temperature

Holding temperature constant, we examined variability in epidemic dynamics across four temperatures: 20°C, 25°C, 30°C, and 35°C. As temperature increased from 20°C to 30°C, the number of susceptible individuals depleted more rapidly (Fig. 2, *S_H_*). At 20°C and 35°C, the epidemics were small (1.33% and 5.92% of the population infected, respectively) and burned out rapidly. Although simulations run at 25°C and 30°C produced final epidemic sizes of 94.73% and 99.98% of the population infected, respectively (Fig. 2, *R_H_*), the epidemic peaked much faster at 30°C.

**Fig 2.**
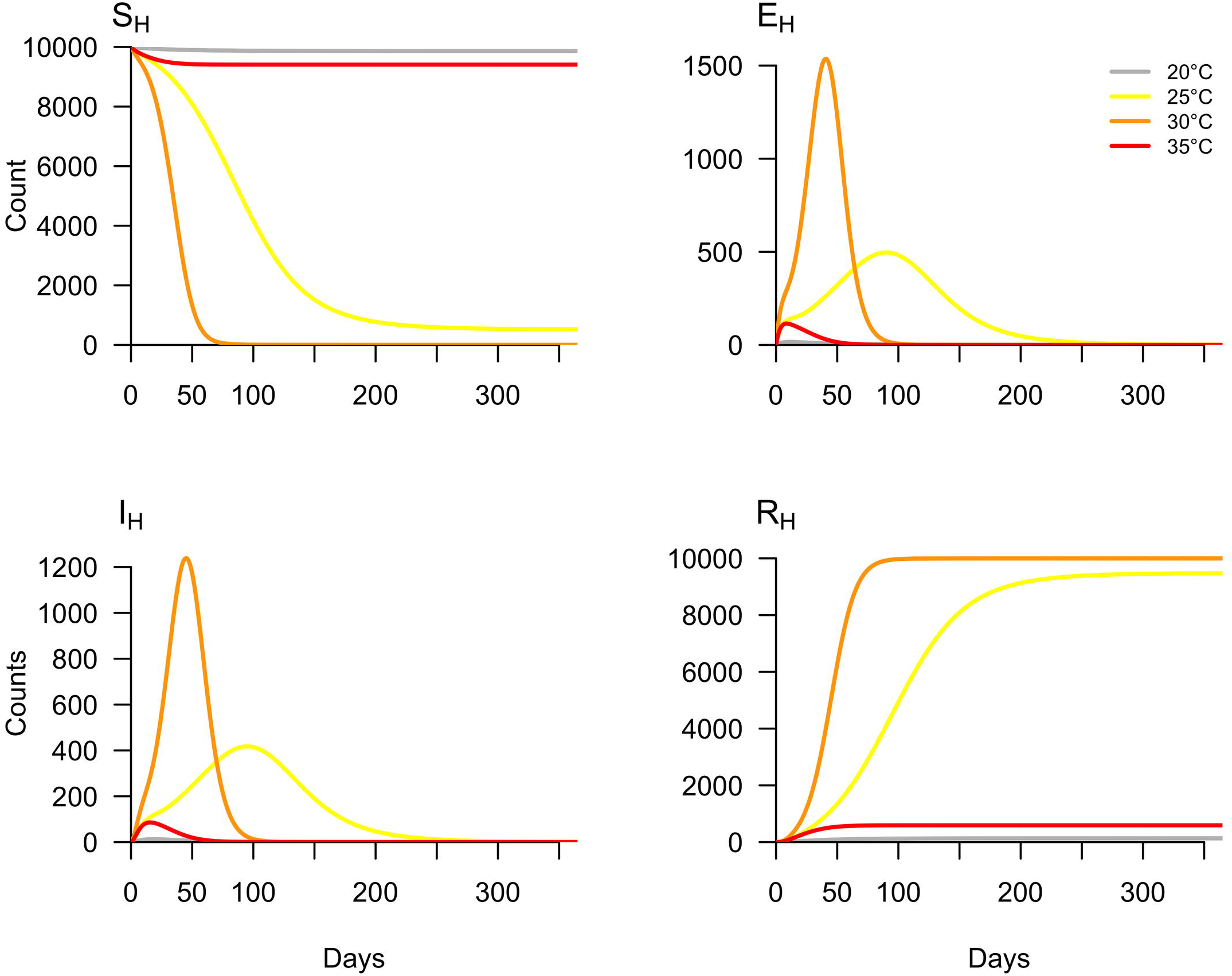
Variation in epidemic dynamics by temperature. The model was simulated under default parameters at four constant temperatures: 20°C, 25°C, 30°C, and 35°C.

### Variability in Epidemic Dynamics with Starting Temperature

Next, we examined variability in epidemic dynamics due to the temperature at which the epidemic began, given two seasonal temperature regimes (25°C mean and a seasonal range of 10°C to 40°C or 20°C to 30°C, respectively). Given that an epidemic occurred, epidemic length monotonically decreased as a function of starting temperature for the first temperature regime (Fig. 3, A): warmer temperatures at the start of the epidemic produced shorter epidemics, and vice versa. In the second temperature regime, epidemic length monotonically decreased as a function of starting temperature until ~29°C. When temperature varied from 10°C to 40°C, the longest epidemic simulated was 137.8 days and occurred at starting temperatures of 11.2°C, and the shortest epidemic lasted 16.82 days and occurred when the temperature at the epidemic start was 35.7°C. When the temperature was 35.8°C or higher or 10.2°C or lower, no epidemic occurred. When temperature was constrained between 20°C and 30°C, the longest epidemic simulated was 253.64 days at a starting temperature of 20°C, and the shortest epidemic lasted 136.1 days at a starting temperature of 28.9°C.

**Fig 3.**
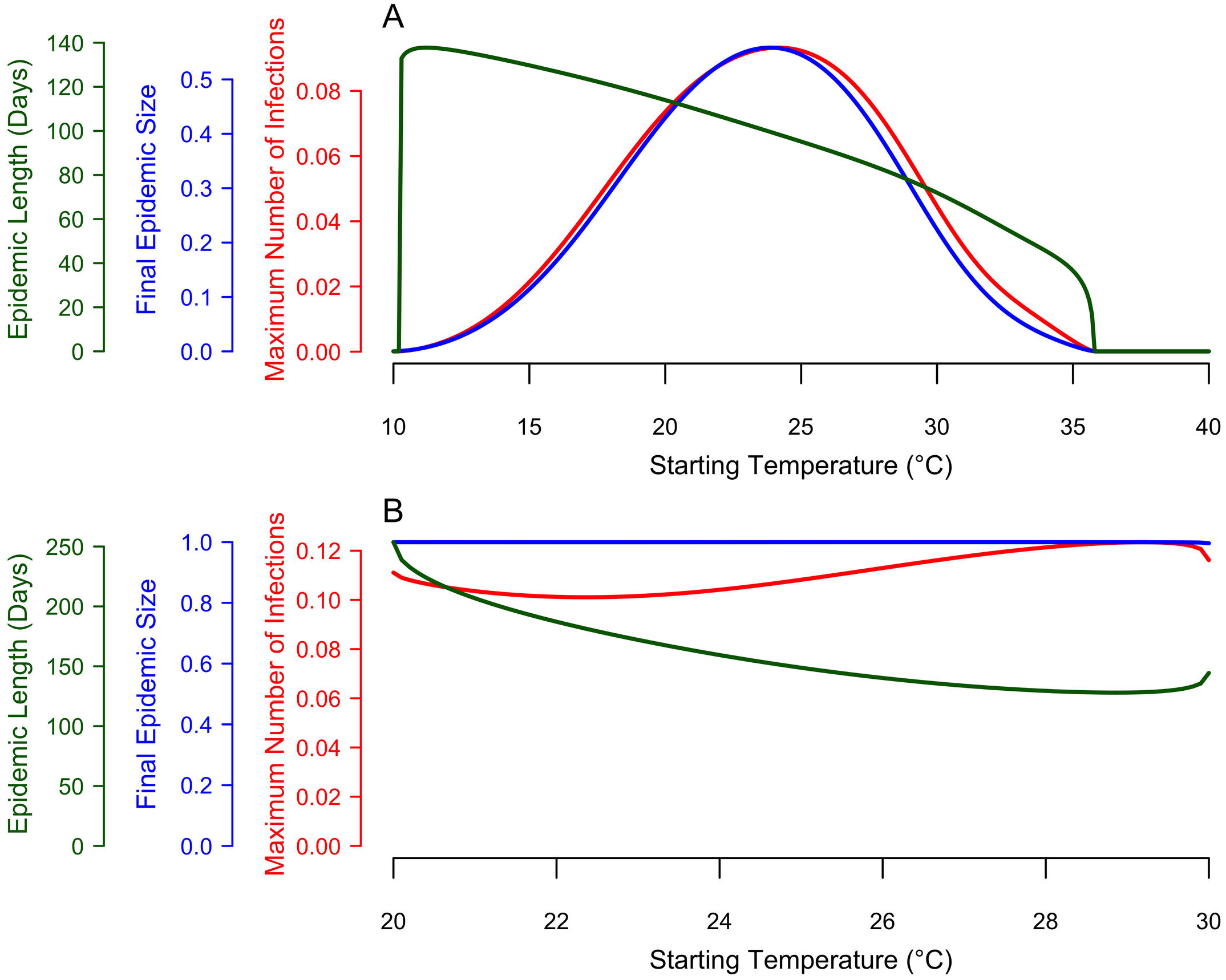
Epidemiological indices as a function of starting temperature, within a given seasonal temperature regime. The red curve represents the maximum number of humans in the infected class (*I_H_*) at any given point during the simulation. The blue curve represents the final (or cumulative) epidemic size (*R_H_* at the final time step). The green curve represents the length of the epidemic (i.e., the point at which the number of infected individuals was below one). Here, simulations were run with the temperature conditions: *T_min_* = 10°C, *T_mean_* = 25°C, and *T_max_* = 40°C (A) and *T_min_* = 20°C, *T_mean_* = 25°C, and *T_max_* = 30°C (B).

In contrast to epidemic length, the response of final epidemic size and maximum number of infected individuals to the temperature at epidemic onset depended on the amount of seasonal temperature variation. When temperature varied widely, from 10°C to 40°C, both final epidemic size and the maximum number of infected individuals responded unimodally to starting temperature, with peaks at 23.9°C and 24.1°C, respectively (Fig. 3, A). By contrast, when temperature varied more narrowly from 20°C to 30°C, the final epidemic size and the maximum number of infected individuals were insensitive to starting temperature (Fig. 3, B). Taken together, these results show that epidemics introduced at different times within identical seasonal temperature regimes can produce very similar final epidemic sizes and maximum infection rates, provided that the temperature range is sufficiently constrained. If temperature variation is large, dramatically different final epidemic sizes and maximum infection rates may result.

### Seasonal Variability of Final Epidemic Size

To address how mean temperature and seasonal variance combined to influence the final epidemic size, we simulated over a wide range of temperature regimes that accounted for variation in the mean and temperature range over a calendar year.

We calculated relative epidemic suitability, defined as the final epidemic size as a proportion of the human population, for twenty major cities worldwide (Table 3).

In a low-variation thermal environment, a band of mean temperatures between approximately 25°C and 35°C supports the highest epidemic suitability (Fig. 4). As the seasonal temperature range increases, lower mean temperatures are capable of supporting large epidemics. However, outside this narrow band of temperature regimes, epidemic suitability rapidly diminishes, and most temperature regimes did not produce epidemics.

**Fig 4.**
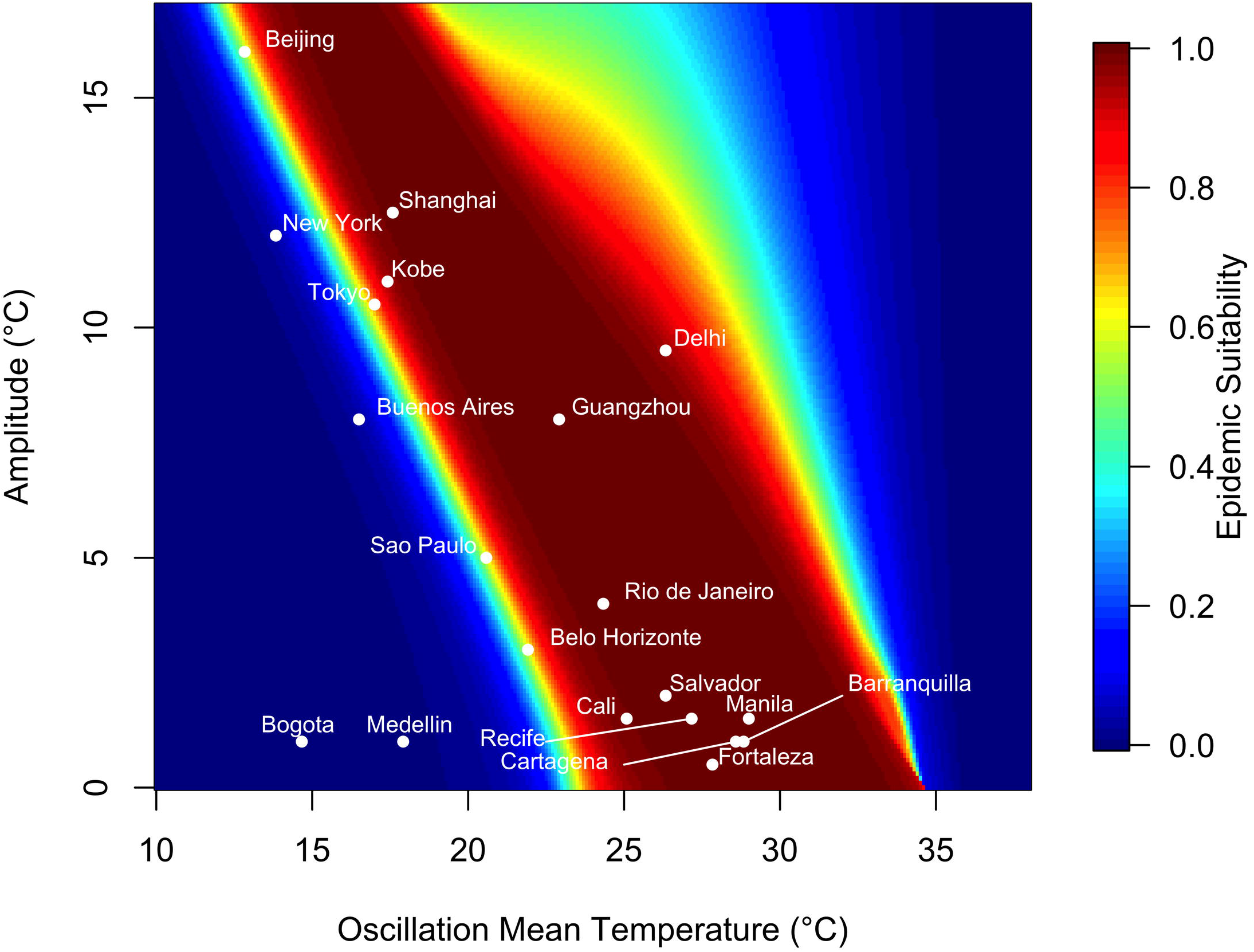
Variation in epidemic suitability across different seasonal temperature regimes. The heat map shows the epidemic suitability (represented as the proportion of the total human population infected during an epidemic) as a function of mean annual temperature and temperature range. Here, temperature range is defined as the seasonal variation about the annual mean temperature. Twenty large, globally important cities are plotted to illustrate their epidemic suitability.

Of the focal 20 major cities, those with high mean temperature and small average temperature variation exhibited the highest epidemic suitability. For instance, Manila, Philippines, which has a monthly mean temperature of 29°C and average seasonal amplitude in mean temperature of 1.50°C, had an epidemic suitability of 0.9998. Cartagena and Barranquilla, Colombia had epidemic suitability of 0.9997. On the other hand, areas with low average temperature and greater temperature variation, such as Beijing and New York, exhibited lower—but still non-zero—epidemic suitabilities of 0.5268 and 0.04088 respectively. Notably, Guangzhou and Shanghai, China have high epidemic suitability (0.9996 and 0.9966, respectively) despite moderate mean temperatures (22.9 and 17.6°C, respectively) due to high seasonal variation in temperature. By contrast, high seasonal variation reduced suitability to 0.9537 in Delhi, India, which has a high mean temperature of 26.3°C (Fig. 4).

The relationship between epidemic suitability and seasonal temperature regime was consistent across varying levels of population immunity. Locations with high mean temperatures and small average temperature variation had higher epidemic suitability, regardless of the level of population immunity (Figures S2-S5). However, as the level of immunity increased from 20% to 80%, the epidemic suitability at given seasonal temperature regime decreased (Table 3).

Epidemic suitability also varied by starting temperature, depending on the seasonal temperature regime. The epidemic suitability of cities with high mean temperature and small average temperature variation—such as Manila, Philippines and Cartagena and Barranquilla, Colombia—did not depend on starting temperature (Table 4). However, areas with low to moderate mean temperature and large average temperature variation (e.g., Kobe, Japan and Shanghai, China) exhibited low epidemic suitability (both 0.0001000) at the minimum starting temperature and moderate-to-high epidemic suitability at the maximum starting temperature (0.6890 and 0.8905, respectively) (Fig. 5). The opposite occurred in regimes with high mean temperature and large temperature variation, though these temperature regimes are rarer.

**Fig 5.**
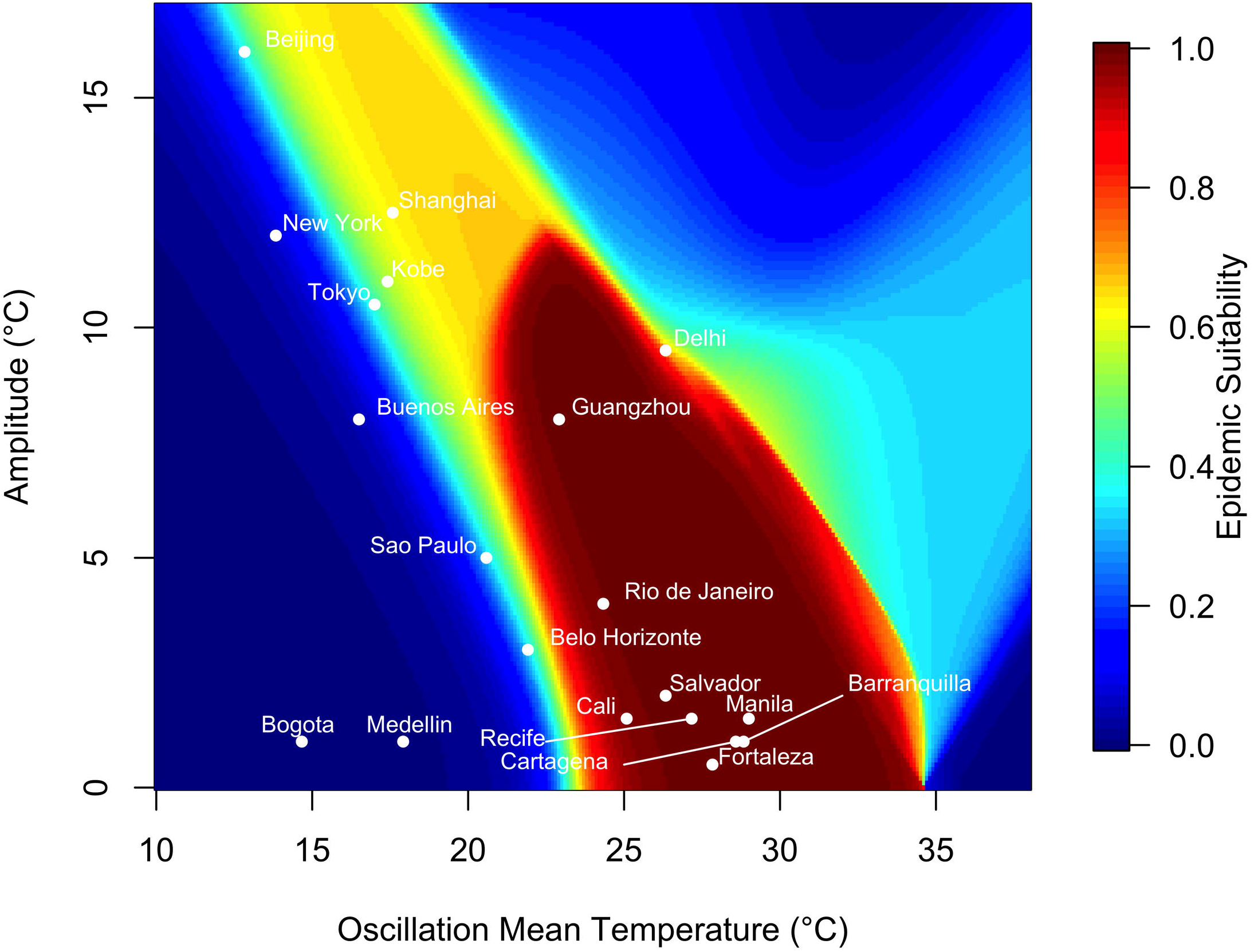
Variation in epidemic suitability across different seasonal temperature regimes averaged across starting temperatures. The heat map shows the epidemic suitability (represented as the proportion of the total human population infected during an epidemic) as a function of mean annual temperature and temperature range averaged across simulations where the initial temperature was set to the seasonal temperature regime’s minimum, mean, or maximum temperature. Here, temperature range is defined as the seasonal variation about the annual mean temperature. Twenty large, globally important cities are plotted to illustrate their epidemic suitability.

**Table 4.** Estimates of epidemic suitability for major cities under different starting temperatures. Epidemic suitability was calculated as the proportion of the population that became infected in simulations that began at the minimum, mean, or maximum temperature of the seasonal temperature regime. Each city was simulated with its respective temperature regime from the 2016 calendar year with 0% population immunity.

Epidemic Suitability
| City | Minimum Starting<br>Temperature | Mean Starting<br>Temperature | Maximum Starting<br>Temperature |
| --- | --- | --- | --- |
| Buenos Aires, Argentina | 0.0001000 | 0.03656 | 0.1166 |
| Sao Paulo, Brazil | 0.02026 | 0.6056 | 0.3480 |
| Rio de Janeiro, Brazil | 0.9978 | 0.9984 | 0.9760 |
| Salvador, Brazil | 0.9965 | 0.9990 | 0.9963 |
| Fortaleza, Brazil | 0.9986 | 0.9993 | 0.9990 |
| Belo Horizonte, Brazil | 0.09404 | 0.5909 | 0.3273 |
| Recife, Brazil | 0.9973 | 0.9994 | 0.9987 |
| Shanghai, China | 0.0001000 | 0.9966 | 0.8905 |
| Beijing, China | 0.0001000 | 0.5268 | 0.5792 |
| Guangzhou, China | 0.9983 | 0.9996 | 0.9912 |
| Bogotá, Colombia | 0.0001000 | 0.0001000 | 0.0001000 |
| Medellin, Colombia | 0.0002177 | 0.002544 | 0.004472 |
| Cali, Colombia | 0.9858 | 0.9909 | 0.9623 |
| Barranquilla, Colombia | 0.9994 | 0.9997 | 0.9997 |
| Cartagena, Colombia | 0.9993 | 0.9997 | 0.9997 |
| Delhi, India | 0.5615 | 0.9537 | 0.6954 |
| Tokyo, Japan | 0.0001000 | 0.7269 | 0.5121 |
| Kobe, Japan | 0.0001000 | 0.9435 | 0.6890 |
| Manila, Philippines | 0.9994 | 0.9998 | 0.9998 |
| New York, USA | 0.0001000 | 0.04088 | 0.1863 |

Estimated epidemic suitability is close to one in the most suitable temperature regimes because we assumed that: (i) the population was fully susceptible at the start of the epidemic; (ii) mixing was homogeneous among humans and mosquitoes; (iii) all cases of infection are included regardless of whether or not they are symptomatic; and (iv) no other environmental or social drivers are limiting transmission. As a result, the epidemic suitability metric should be considered an upper bound on the proportion of the population that could become infected based on temperature alone.

### Model Sensitivity and Uncertainty Analysis

Final epidemic size was not sensitive to life history trait parameterization (Figs. S8-S10), using samples from the posterior distribution of thermal response fits for each temperature-dependent trait.

There was uncertainty in the specific numerical values of the epidemiological indices across starting temperatures (Fig. S1). However, the overall functional response of the final epidemic size, maximum number of infected individuals, and the epidemic length to starting temperature was consistent across the samples from the joint posterior distribution.

## Discussion

Recent outbreaks of DENV, CHIKV, and ZIKV in Latin America and across the globe have captured the attention of the public health community and underscore the importance of preparation for future outbreaks. As temperatures rise, the global landscape suitable for such outbreaks will expand and shift geographically, potentially placing a larger proportion of the world’s population at risk [24,29,31]. Understanding how local temperature regimes govern epidemic dynamics is increasingly important for determining resource allocation and control interventions [41]. While previous work has investigated the effects of temperature on DENV, CHIKV, and/or ZIKV transmission, until now we have lacked comprehensive, mechanistic, and dynamic understanding of the effects of seasonally varying temperature on transmission via its (nonlinear) effects on mosquito and parasite traits [27–34]. With our model, which expands on [24] and [25], we show that seasonal temperature mean and amplitude interact with the temperature at epidemic onset to shape the speed and magnitude of epidemics.

At constant temperature, epidemics varied substantially in the rate at which susceptible individuals were depleted. Epidemics simulated at 25°C and 30°C reached similar sizes but the epidemic at 25°C proceeded at a much slower rate (Fig. 2). This “slow burn” phenomenon occurs because slower depletion of susceptible individuals can produce epidemics of similar size to epidemics that infect people very rapidly. This phenomenon also occurs in more realistic, seasonally varying temperature regimes.

The temperature at which an epidemic started affected dynamics only under large ranges of temperature variation. When temperature ranged from 10°C to 40°C, the final epidemic size peaked at intermediate starting temperatures (24°C; Fig. 3, A). However, in highly suitable seasonal environments, final epidemic size was large regardless of the starting temperature (Fig. 3, B).

At mean starting temperatures, epidemic suitability was sensitive to the interaction between annual temperature mean and seasonal variation. Under low seasonal temperature variation, a narrow band of annual mean temperatures (approximately 25-35°C) had the highest epidemic suitability (Figs. 4 & S2-S5). Outside this band of temperature regimes, suitability diminishes rapidly. Larger seasonal variation in temperature lowers the range of optimal annual mean temperatures (i.e., suitability is high in cooler places with larger seasonal variation in temperature; Fig. 4).

The relationship between epidemic suitability and the seasonal temperature regime also depended on the temperature at the epidemic onset. Three distinct relationships emerged (Figs. 5 & S6-S7). At intermediate annual mean temperatures of ~25-35°C and low seasonal temperature variation (~0-10°C), epidemic suitability is insensitive to starting temperature because temperature is suitable for transmission year-round. At lower annual mean temperatures (~10-25°C) and higher seasonal temperature variation (~10-15°C), epidemic suitability is highest when epidemics start in moderate to warm seasons, and lower when epidemics start during cooler seasons. Finally, at high annual mean temperatures (> 35°C) and low seasonal temperature variation (~0-10°C), epidemic suitability is high only when epidemics start at the coldest period of the year, because otherwise the temperature is too warm for efficient transmission. The interaction between temperature mean, annual variation, and starting point sharply illustrates the unimodal effect of temperature on transmission. Models that do not include unimodal effects of temperature (e.g., those with sinusoidal forcing on a transmission parameter) may fail to capture the limits on transmission in warm environments.

With rising mean annual temperatures and increasing seasonal temperature variation due to climate change, the landscape of epidemic suitability is likely to shift. Importantly, areas with previously low epidemic suitability may have increasing potential for transmission year-round. By contrast, warming temperatures may drive epidemics in cities with high current suitability (e.g., Manila, Philippines, Barranquilla, Colombia, and Fortaleza, Brazil) to shift toward cooler months. Thus, climate change may alter not only epidemic size and duration but also seasonal timing globally, as it interacts with other important drivers like rainfall and human behavior.

It is important to note that model-estimated epidemic suitability should be treated as an upper bound on the potential for large epidemics because within highly suitable climate regimes, epidemics can vary in magnitude due to human population size and movement dynamics [28], effective vector control, and other mitigating factors. Likewise, our estimates are conditioned on *Aedes aegypti* presence and virus introduction to support an outbreak.

Although seasonal temperature dynamics provide insight into vector-borne transmission dynamics, other factors like mosquito abundance, vector control, and rainfall also determine transmission dynamics. Thus, temperature must be considered jointly with these factors. Moreover, accurately describing epidemic dynamics of emerging and established vector-borne pathogens will ultimately require integrating realistic models of environmental suitability, as presented here, with demographic, social, and economic factors that promote or limit disease transmission [42,43]. Conversely, we show that the interaction between temperature and the availability of susceptible hosts alone can drive epidemic burnout even in the absence of other limiting factors like vector control and seasonal precipitation. This suggests that correctly representing the nonlinear relationship between temperature and epidemic dynamics is critical for accurately inferring mechanistic drivers of epidemics and, in turn, predicting the efficacy of control interventions.

## Supporting Information Legends

**S1 Fig. Sensitivity of epidemiological indices as a function of starting temperature to the parametrization of life history traits**. The red curve represents the median maximum number of humans in the infected class (*I_H_*) at any given point during the simulation. The blue curve represents the median final (or cumulative) epidemic size (*R_H_* at the final time step). The green curve represents the median length of the epidemic (i.e., the point at which the number of infected individuals was below one). Each shaded area represents the 95% credible interval for the epidemiological indices ran under 50 different parameterizations of the life history traits. Here, simulations were run with the temperature conditions: *T_min_* = 10°C, *T_mean_* = 25°C, and *T_max_* = 40°C (A) and *T_min_* = 20°C, *T_mean_* = 25°C, and *T_max_* = 30°C (B).

**S2 Fig. Variation in epidemic suitability across different seasonal temperature regimes with 20% population immunity**. The heat map shows the epidemic suitability (represented as the proportion of the total human population infected during an epidemic) as a function of mean annual temperature and temperature range assuming 20% population immunity. Here, temperature range is defined as the seasonal variation about the annual mean temperature. Twenty large, globally important cities are plotted to illustrate their epidemic suitability.

**S3 Fig. Variation in epidemic suitability across different seasonal temperature regimes with 40% population immunity**. The heat map shows the epidemic suitability (represented as the proportion of the total human population infected during an epidemic) as a function of mean annual temperature and temperature range assuming 40% population immunity. Here, temperature range is defined as the seasonal variation about the annual mean temperature. Twenty large, globally important cities are plotted to illustrate their epidemic suitability.

**S4 Fig. Variation in epidemic suitability across different seasonal temperature regimes with 60% population immunity**. The heat map shows the epidemic suitability (represented as the proportion of the total human population infected during an epidemic) as a function of mean annual temperature and temperature range assuming 60% population immunity. Here, temperature range is defined as the seasonal variation about the annual mean temperature. Twenty large, globally important cities are plotted to illustrate their epidemic suitability.

**S5 Fig. Variation in epidemic suitability across different seasonal temperature regimes with 80% population immunity**. The heat map shows the epidemic suitability (represented as the proportion of the total human population infected during an epidemic) as a function of mean annual temperature and temperature range assuming 80% population immunity. Here, temperature range is defined as the seasonal variation about the annual mean temperature. Twenty large, globally important cities are plotted to illustrate their epidemic suitability.

**S6 Fig. Variation in epidemic suitability across different seasonal temperature regimes with minimum starting temperature**. The heat map shows the epidemic suitability (represented as the proportion of the total human population infected during an epidemic) as a function of mean annual temperature and temperature range. Here, temperature range is defined as the seasonal variation about the annual mean temperature, and the simulation began at the minimum temperature of the regime. Twenty large, globally important cities are plotted to illustrate their epidemic suitability.

**S7 Fig. Variation in epidemic suitability across different seasonal temperature regimes with maximum starting temperature**. The heat map shows the epidemic suitability (represented as the proportion of the total human population infected during an epidemic) as a function of mean annual temperature and temperature range. Here, temperature range is defined as the seasonal variation about the annual mean temperature, and the simulation began at the maximum temperature of the regime. Twenty large, globally important cities are plotted to illustrate their epidemic suitability.

**S8 Fig. The 2.5% quantile of epidemic suitability to the parameterization of life history traits**. Epidemic suitability (represented as the proportion of the total human population infected during an epidemic) as a function of mean annual temperature and the temperature range. Temperature varied according to a seasonal temperature regime, and 50 samples of c, Tmin, and Tmax were taken from the joint posterior distribution of each trait thermal response from Mordecai et al. [24].

**S9 Fig. The 50% quantile of epidemic suitability to the parameterization of life history traits**. Epidemic suitability (represented as the proportion of the total human population infected during an epidemic) as mean annual temperature and the temperature range. Temperature varied according to a seasonal temperature regime, and 50 samples of c, Tmin, and Tmax were taken from the joint posterior distribution of each trait thermal response from Mordecai et al. [24].

**S10 Fig. The 97.5% quantile of epidemic suitability to the parameterization of life history traits**. Epidemic suitability (represented as the proportion of the total human population infected during an epidemic) as mean annual temperature and the temperature range. Temperature varied according to a seasonal temperature regime, and 50 samples of c, Tmin, and Tmax were taken from the joint posterior distribution of each trait thermal response from Mordecai et al. [24].

